# Spontaneous behavioral coordination between pedestrians emerges through mutual anticipation rather than mutual gaze

**DOI:** 10.1101/2022.07.10.499066

**Authors:** Hisashi Murakami, Takenori Tomaru, Claudio Feliciani, Yuta Nishiyama

**Affiliations:** Faculty of Information and Human Science, Kyoto Institute of Technology, Kyoto-shi, Kyoto, Japan; Research Center for Advanced Science and Technology, The University of Tokyo, Meguro-ku, Tokyo, Japan; Information and Management Systems Engineering, Nagaoka University of Technology, Nagaoka, Niigata, Japan

**Keywords:** coordination dynamics, pedestrian interaction, human crowd, anticipation, distraction

## Abstract

Human activities are often performed together between two or more persons, as if they are a complex dance. Threading through a crowd is a striking example of such coordinated actions. Behavioral coordination should help to reduce head-on collisions, smooth a pedestrian’s pathway through a crowd, and promote a self-organization process. Although mutual anticipation between pedestrians would be a candidate for underlying mechanisms of behavioral coordination, it remains largely unexplored, especially in terms of visual information. Here, we investigated the effects of mutual anticipation between a pair of pedestrians performing simple avoidance tasks using a combination of motion- and eye-tracking systems. We found that pedestrians in a baseline condition spontaneously coordinated their walking speed and angle until passing each other. Visually distracting one of the pedestrians decreased the level of behavioral coordination, indicating that spontaneous coordination emerges through mutual anticipation. Importantly, blocking the pedestrians’ gaze information alone did not alter their walking, clearly ruling out the assumption that mutual gaze impacts pedestrian anticipation behavior. Moreover, eye-movement analysis demonstrated that the direction of a pedestrian’s gaze changed depending on the uncertainty of the oncoming pedestrian’s motion and that pedestrians tend to look ahead toward the ultimate passing direction before they actually walked in that direction. We propose that body motion cues may be sufficient and available for implicit negotiation on potential future motions. Our findings should be useful in various fields, including research on improvisational motions, pedestrian transportation, and robotic navigation.

## Introduction

Human activities are often performed together between two or more persons, as if they are a complex dance [1, 2]. Even in day-to-day experiences of non-experts of physical expression, there is an astonishingly rich variety of coordinated human behaviors. Such behaviors can be found in a wide range of social interactions [3-6], from concrete motor actions, like finger tapping and Mexican waves through a large crowd, to abstract social processes, such as engaging in conversation and building consensus. To coordinate with others, people must continuously anticipate the moves of others [1].

Threading through a crowded street is a striking example of these coordinated actions taken among strangers who anticipate each other’s movements. Indeed, although pedestrians have been assumed to be passively repelled by others to avoid collisions in classical theoretical models [7-9], recent empirical and theoretical studies emphasize that pedestrians’ interactions are fundamentally anticipatory in nature [10-16]. In other words, their motions are essentially influenced by the anticipated future positions of neighbors rather than their current positions. Moreover, “mutual” anticipation between pedestrians enables successful collision avoidance and thereby facilitates efficient pattern formation in human crowds, such as the spontaneous creation of unidirectional lanes in bidirectional flow [13, 14]. However, if there are distracted walkers who are unable to anticipate others’ motions in a pedestrian flow, the non-distracted pedestrians also have difficulties avoiding collisions, and thereby the self-organization process is disturbed through the propagation of reduced levels of anticipation in the crowd [14]. Understanding anticipatory interactions among pedestrians is, therefore, important to help manage mass events and evacuation strategies and to design daily pedestrian transportation. However, the underlying mechanisms of behavioral coordination and the visual information required to anticipate have remained largely uninvestigated.

If two interacting pedestrians mutually anticipate each other’s movements and perform a real-time negotiation where both action planning and its execution can be performed simultaneously to avoid collision [14, 17], they would show coordinated joint actions depending on the motions of the other oncoming pedestrian. However, to what extent they coordinate their actions remains poorly understood. Moreover, in the context of social interactions, gaze has been considered to be one of several core processes, for example, the concept of mutual gaze [18, 19]. However, in the case of pedestrian interactions, the idea that pedestrians infer other people’s movement trajectories from the gaze direction remains controversial [20, 21]. Contributing to this controversy may be the fact that experiments on this topic have typically been carried out in a virtual reality setup. Although the few real pedestrian experiments have been conducted using a wearable eye-tracking system [22, 23], a combination of eye- and motion-tracking systems while walking is necessary to measure when and where pedestrians look at other people. In particular, where and to what extent pedestrians look at other pedestrians to anticipate their movements has been largely unexplored. Although additional investigations are needed to clarify the coordination processes that are accompanied by eye movements, existing crowd experiments are inadequate because it is difficult to distinguish effects from multiple neighbors.

This study investigated the effects of mutual anticipation between pairs of pedestrians by means of a combination of motion- and eye-tracking systems (see Materials and methods; Fig. 1A). We conducted simple avoidance task experiments where two pedestrians walked toward and passed each other along a mock corridor. Because of its elementary form of interaction, this type of passing behavior by two pedestrians is one of the most fundamental situations that enables us to directly quantify the behavioral effects of mutual anticipation. We set three experimental conditions: baseline (BASE, namely mutual anticipation is available), no mutual anticipation (NMA), and no mutual gaze (NMG). One of the two participants (represented by P2) wore eye-tracking glasses to measure their gaze behavior, and the other (represented by P1) was asked to do a specific task in accordance with experimental conditions. In NMA, P1 performed a typing task requiring concentration on a mobile phone while walking. This task has been shown to interfere with pedestrians’ ability to anticipate neighbors’ motions [14] because the use of a mobile phone while walking narrows the user’s perceptual visual field [24] and distracts visual attention [25]. In NMG, P1 wore mirrored sunglasses, which blocked the transfer of gaze information from P1 to P2. In the BASE condition, P1 was asked to do nothing special, and the two participants just walked through the corridor.

**Figure 1.**
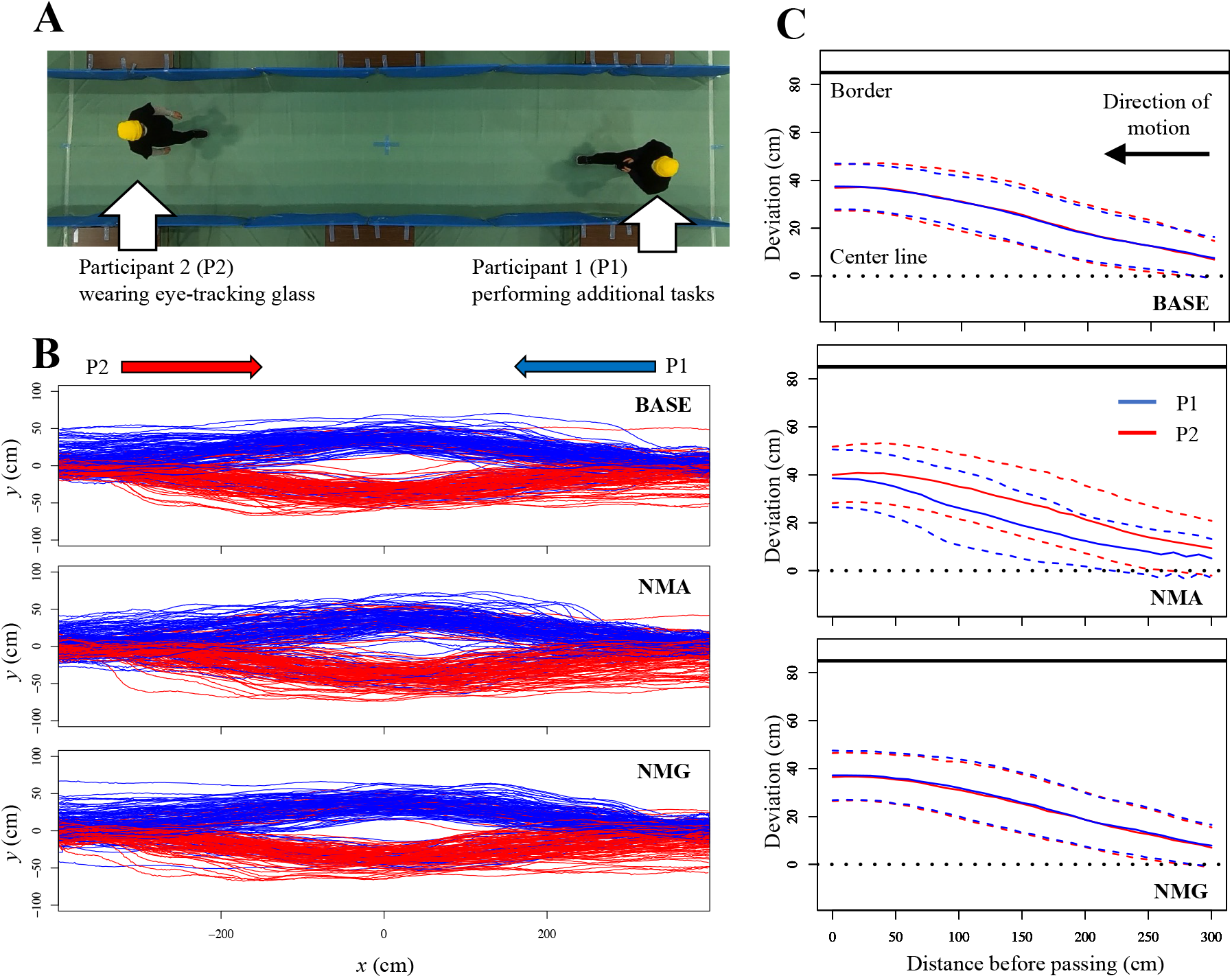
Simple avoidance task experiments manipulating mutual anticipation and mutual gaze. (**A**) A photo from an experiment under the NMA condition. One of two participant pedestrians (P1) performs an additional task when walking (using a smart phone and wearing sunglasses under the NMA and NMG conditions, respectively; no additional task under the BASE condition), while the other pedestrian (P2) wears eye-tracking glasses. BASE, baseline condition; NMA, no mutual anticipation condition; NMG, no mutual gaze condition. (**B**) Reconstructed pedestrian trajectories under each condition. Blue (red) lines represent P1s (P2s) directed to move from left (right) to right (left). (**C**) Average lateral deviation before passing under each condition. Dashed lines represent the standard deviation.

We observed the behavioral coordination between a pair of pedestrians engaging in mutual anticipation. They continuously adjusted their motions relative to each other so that walking speed and direction changed together under the BASE condition. In other words, pedestrians that were about to pass each other did not necessarily follow their own predetermined path, making blind assumptions about oncoming person’s movements. This coordination decreased under NMA, but not under NMG, clearly ruling out the assumption that mutual gaze impacts pedestrian anticipation behavior. In addition, results from the eye-tracking systems not only demonstrated how pedestrians change their gaze behavior depending on their uncertainty about the other walker, but also revealed that they looked ahead toward the future ultimate passing direction before they actually walked in that direction.

## Results

### Influence of mutual anticipation and mutual gaze on walk trajectories

Two pedestrians initially located at the center of their respective starting lines began to walk toward each other (i.e., in opposite directions) along the experimental corridor at the same time. They would deviate from the center line to avoid each other in advance, then pass each other, and finally arrive at their respective destinations (Fig. 1A and 1B). There was a difference in walk trajectories between conditions. The pair of pedestrians (P1 and P2) seemed to deviate in a similar way in the BASE and NMG conditions, but deviated in a different way in NMA. To clarify this, we calculated the absolute lateral deviation along the *y*-axis (perpendicular to the corridor length) as a function of the distance along the *x*-axis from the future *x*-position at which the pedestrians would pass each other, following the procedure used in previous studies [26, 27]. The distance ranged from 0 to 300 cm and was divided into 30 bins, each of which represented 10 cm.

In the BASE condition, the average deviations of P1 and P2 had a similar pattern (Fig. 1C). In contrast, in NMA, there was a clear difference between the two trajectories. The deviation of P2 was greater than that in the BASE condition: P1 initially walked near the center line and then made a large turn immediately before passing (see also Fig. S1, which presents comparisons of the trajectories between the BASE and NMA conditions). In addition, we found that distance between the two at the moment of passing in NMA was larger than that in the BASE condition (Fig. 2A; see Table S1), indicating that pedestrians in NMA tried to keep a larger distance between themselves than they did in the BASE condition. These results imply that the visual distraction induced by the mobile phone activity directly reduced the distracted pedestrian’s anticipation and indirectly altered the non-distracted pedestrian’s behavior as well. These results are in agreement with those of a previous study based on crowd experiments [14].

**Figure 2.**
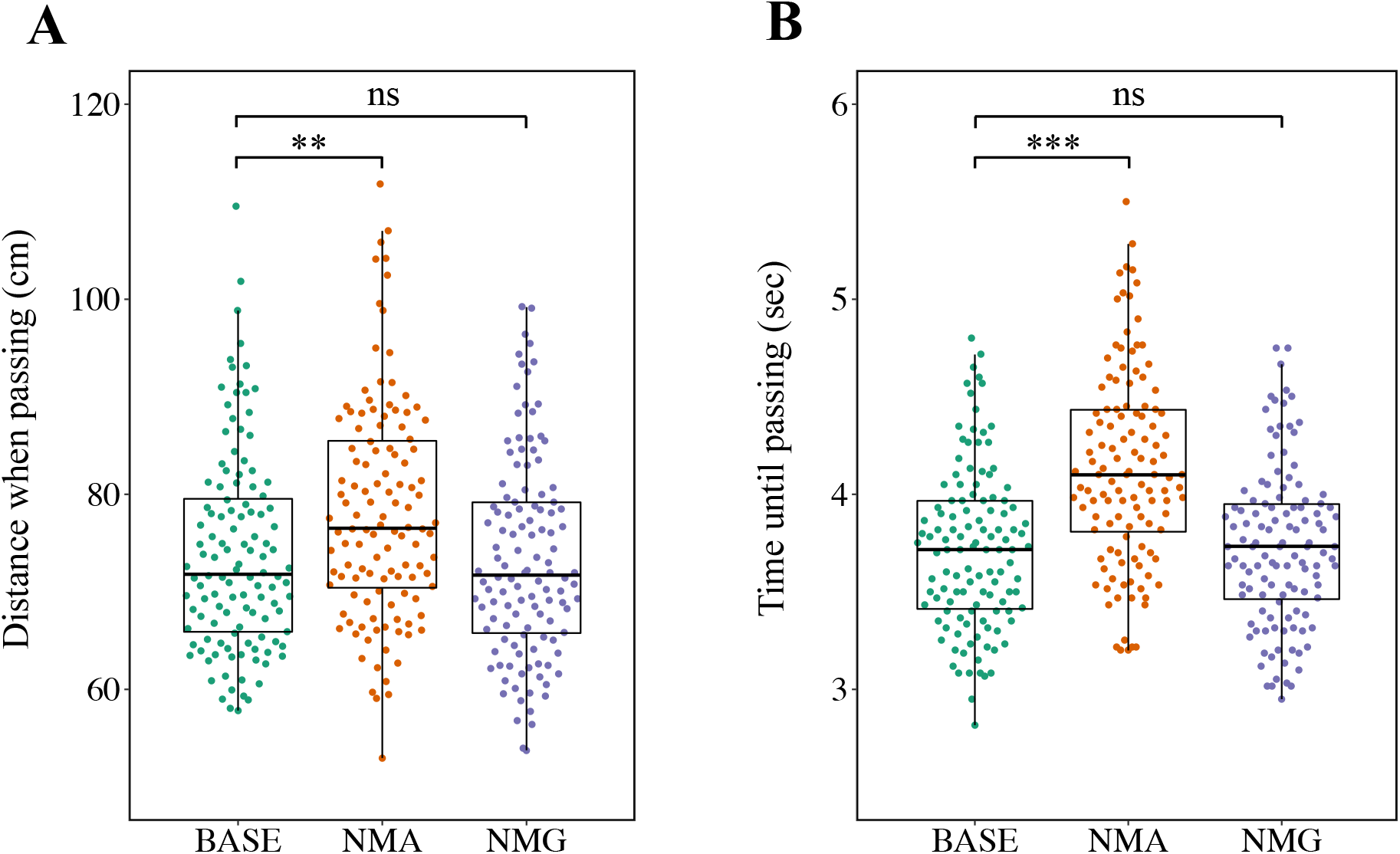
Distance when passing and time until passing. (**A**) Distance between two pedestrians at the moment they pass each other. (**B**) Time taken from the trial onset to the moment of passing. BASE, baseline condition; NMA, no mutual anticipation condition; NMG, no mutual gaze condition. Each data point represents a trial. The statistical comparison is between the BASE condition and each of the other conditions (***p < 0.001; **p < 0.01; ns, p > 0.05). Box-and-whisker plots represent the median (central thick line), the first and third quartiles (box), and 1.5× the interquartile range of the median (whiskers).

When pedestrians faced distracted oncoming pedestrians, they did not coordinate their motions. This may result from absence of gaze information provided by the distracted walkers. With the NMG condition, we could investigate whether the disturbed mutual gaze *per se* would influence pedestrian behavior. Figure 1C shows that the average deviations of P1 and P2 in NMG matched each other well, similar to the BASE condition results. In addition, when comparing the BASE and NMG conditions, the average deviations showed quite similar patterns (Fig. S1). Moreover, we found no significant difference between the BASE and NMG conditions in the distance between P1 and P2 at the moment of passing (Fig. 2A; Table S1). Furthermore, there was no significant difference between the BASE and NMG conditions in the time taken from the trial onset to the moment of passing, although there was between the BASE and NMA conditions (Fig. 2B; Table S1). These results imply that mutual gaze does not influence pedestrians’ behavior, which is in contrast to the findings of previous studies [20]. In other words, pedestrians probably do not use gaze information to infer the future positions of their neighbors.

### Behavioral coordination between interacting pedestrians

As shown in the previous section, in the BASE and NMG conditions, the mean trajectories and their variances seem to be well-matched between two pedestrians (Fig. 1C). To verify whether these similarities result from behavioral coordination, in this section, we focus on the joint actions of the two pedestrians rather than analyzing their motions separately.

To investigate behavioral coordination, we analyzed the relationship of trajectories between actual pairs of pedestrians in comparison with randomly selected pairs (i.e., a random pairing of P1 from one pair and P2 from another pair). We assumed that actual pairs would coordinate their walking speeds and directions if they negotiated their movements with each other. As one indicator, we first determined changes in an individual’s walking speed by calculating differences between two consecutive speeds at time *t* and time *t* − *dt*, where *dt* = 1 (*s*). As another indicator, we determined angular deviations of the velocity vector from the desired direction (i.e., the direct path to their destination: the *x*-axis along the corridor length in this case) by calculating the angle that the velocity vector made with the *x-*axis at time *t*. These indicators were introduced to describe pedestrians’ collision avoidance behavior in a previous study [14].

Figure 3A shows differences in walking speed and angular deviation between P1 and P2 of actual pairs as a function of time; a smaller difference indicates more coordination. Time was normalized to create a relative time axis ranging from 0 (trial onset) to 1 (the moment of passing) (see the Materials and Methods). Although the plots of BASE and NMG show a quite similar pattern for both indicators, those of NMA demonstrate less coordination (Figure 3A). This implies that two interacting pedestrians are coordinated, especially if they are able to use mutual anticipation.

**Figure 3.**
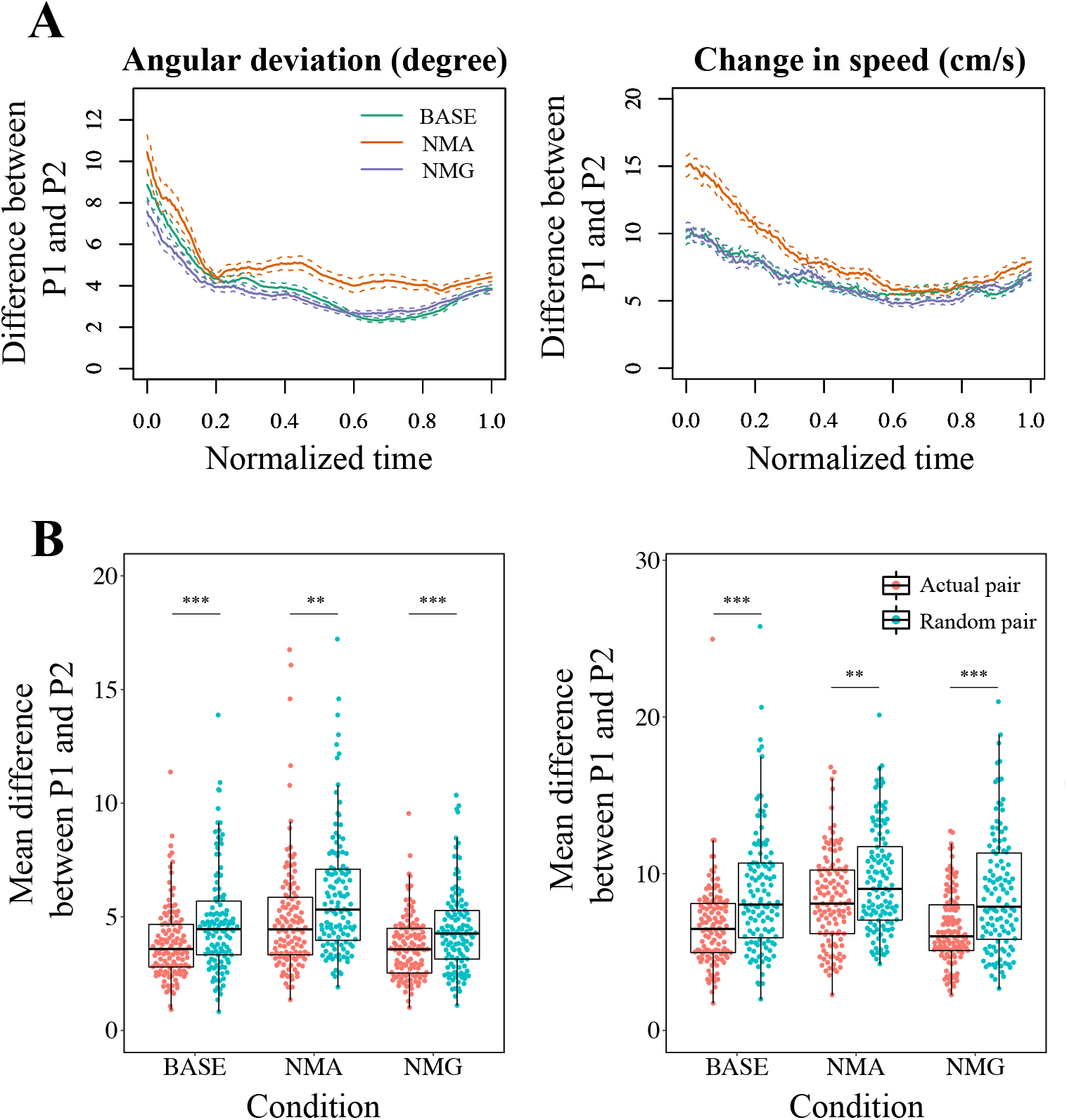
Behavioral coordination between two interacting pedestrians. (**A**) Differences between two interacting pedestrians in angular deviation (left) and change in speed (right) as a function of time. Time was normalized to a relative time axis ranging from 0 (trial onset) to 1 (the moment of passing) as described in the text. The lines show the mean ± SEM (solid and dotted lines, respectively). Both indicators in the NMA condition are always equal to or higher than the other conditions, demonstrating less coordination between the two pedestrians. (**B**) Comparisons between actual and random pairs. Left and right panels show mean differences from the trial onset to the moment of passing for angular deviation and change in speed, respectively. The smaller values in actual pairs than random pairs demonstrate greater coordination of motions between pedestrians who are *actually* interacting. Each data point represents a trial. Asterisks indicate statistical significance (***p < 0.001; **p < 0.01). See Figure 2 for a description of the box-and-whisker plots.

Next, we verified the possibility that two pedestrians simply follow a predetermined path rather than continuously negotiating their motions with each other by including random pairs. We calculated the same time averages for the same indicators and compared the results between actual and random pairs. Note that we calculated the time average from the trial onset according to a shorter data sequence in the random pairs because the time to passing differed between the individuals in the random pairs. In all conditions, the mean differences of actual pairs were smaller than those of random pairs for both angular deviation and change in speed (Figure 3B, see also Table S1). This demonstrates coordination of motions between pedestrians who are *actually* interacting (even in the NMA condition).

### Gaze pattern

In this section, we investigate where and to what extent pedestrians gaze in order to interact with oncoming pedestrians. Because P2 wore eye-tracking glasses equipped with a camera to record the user’s view, we could calculate P2’s gaze points relative to P1 in two dimensions (see Materials and Methods).

To investigate gaze distributions around an oncoming pedestrian according to the distance between P1 and P2 along the *x*-axis, we examined gaze data as a function of the inter-pedestrian *x*-distance, which was divided into 8 sections from 0 cm (the moment of passing) to 800 cm (the maximum distance). Figure 4A shows the two-dimensional distributions of gaze points around oncoming pedestrians at four different sections under the BASE condition (see Figure S2-4 for the other sections and conditions). The human-like silhouette is composed of a circle (centered at the origin [i.e., the head position of P1]) with a radius of 12.5 cm and a rectangle (145 × 46 cm) as the area of interest (AOI) representing the oncoming pedestrian’s body area. The right side of the reference frame in Figure 4A corresponds to the actual passing side. It is easy to see that participants initially looked all over the body of the pedestrian, then gradually shifted their eyes to outside of the AOI in the ultimate passing direction, and finally almost took their gaze off the oncoming pedestrian before passing. To investigate how much the pedestrians look at each other, we calculated the proportion each pedestrian gazed at the AOI in each distance section (see Materials and Methods). In the NMA condition, participants tended to look more at the AOI, namely the body of the oncoming pedestrian, compared to the BASE condition (figure 4B). There were significant differences for half of the distance sections (see Table S2). This stronger visual attention to the AOI under NMA suggests that the mobile-phone-using pedestrians distracted the other pedestrians. On the other hand, the proportions of gaze points on the AOI under NMG were almost the same as those under BASE, except in the 600–700 cm section. We have no explanation for this difference, but it may be related to the results from the follow-up questionnaire survey; in the NMG condition, P2s felt that P1s who wore sunglasses paid less attention to them (see the Materials and Methods, Figure S5, and Table S3). However, please note that this difference in gaze behavior would not influence walking behavior (Figures 1 and 2).

**Figure 4.**
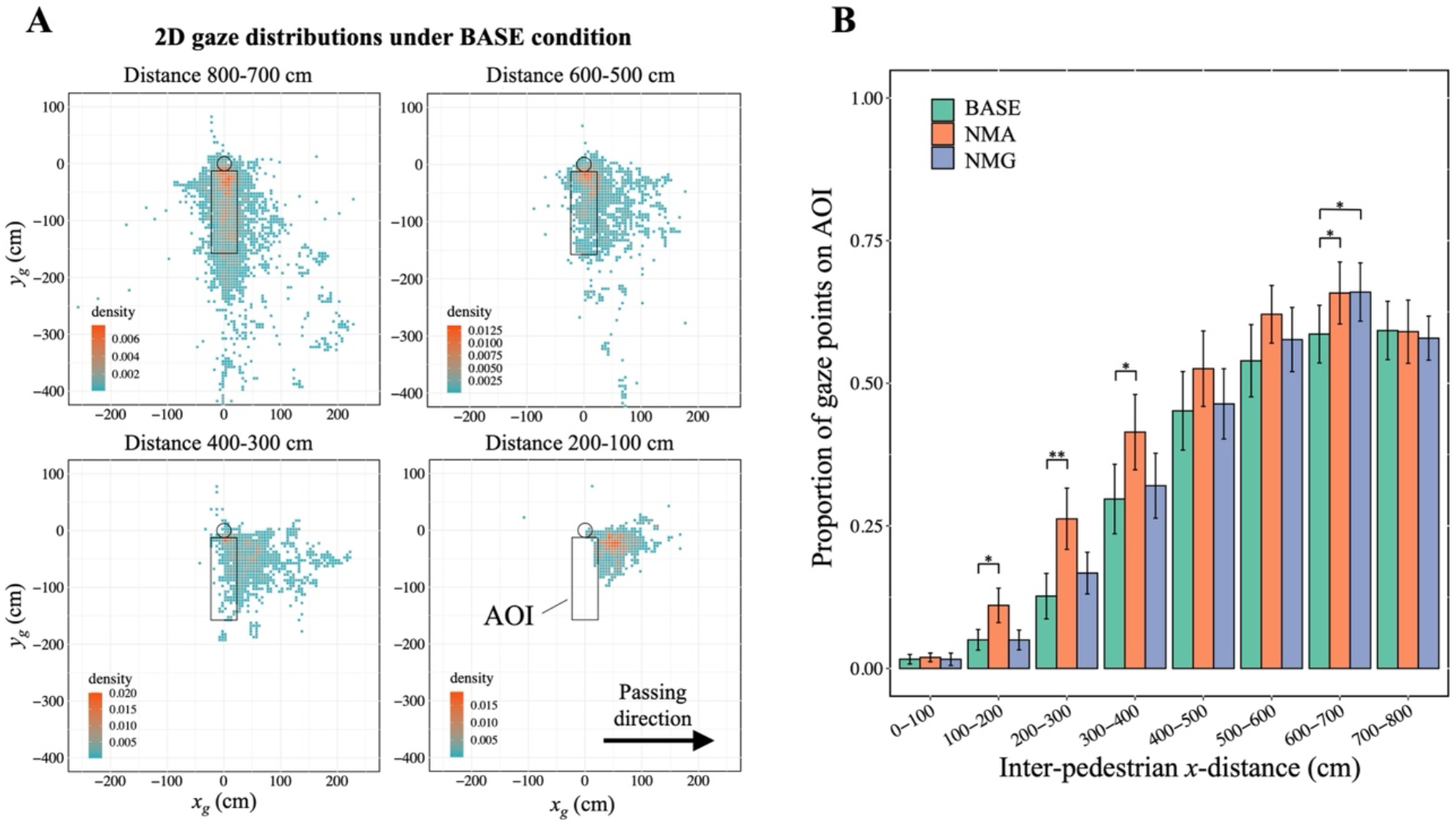
Gaze behavior near oncoming pedestrians while walking. (**A**) Two-dimensional distributions of gaze points around oncoming pedestrians. Four different distance sections between two pedestrians along the *x*-axis under the BASE condition are shown. The human-like silhouette composed of a circle and a rectangle is the area of interest (AOI) representing the oncoming pedestrian’s body area, which was used in the analysis in **B**. *x*_*g*_ and *y*_*g*_ represent horizonal and vertical components of the cm-coordinates of the gaze points relative to P1’s head position, respectively (see the main text). Passing direction was positioned to be right. Bins are 5 × 5 (cm). (**B**) Mean proportion of gaze points on the AOI during sections of inter-pedestrian distance along the *x*-axis. Error bars show SEM. Asterisks indicate statistical significance between the BASE condition and each of the other conditions (**p < 0.01; *p < 0.05).

### Looking ahead toward the ultimate passing direction

We showed that disturbing the mutual gaze has little effect on walk trajectories and behavioral coordination (Figures 1, 2, and 3). The gaze from an oncoming pedestrian is, therefore, unlikely to be a crucial cue in predicting where that person will go. However, this does not necessarily mean that the gaze behavior of pedestrians is unrelated to decision making in terms of their own walking behavior. We therefore investigated the relationship between gaze and which side was chosen for passing. It is well-known that humans tend to gaze at a future moving direction to gain advance information before taking action [28-31]. In our experiments, pedestrians looked all over the body of an oncoming pedestrian at the initial stage of the trial (i.e., distance section 700–800 cm in Figure 4A), probably to gain information to anticipate that person’s behavior. They also gazed at their future passing direction even at the initial stage. To verify this, we calculated the proportion of trials in which participants gazed at the ultimate passing direction for a longer time (than the opposite side) in each distance section (Figure 5A). We found that, even at the initial stage, the proportion was significantly greater than 0.5 (i.e., chance) (Figure 5A inset; Table S3). On the other hand, pedestrians did not necessarily move toward the ultimate passing direction at the initial stage. We calculated the proportion of trials in which participants were on the side of passing at a given inter-pedestrian distance (700, 600, …, 0 cm) on the basis of their *y*-position at the maximum distance (800 cm). We found that the proportion at the initial stage (700 cm in this case) was consistent with chance (Figure 5B; see Table S3). Taken together, these results suggest that participants tend to visually explore their future moving direction even before they move in that direction.

**Figure 5.**
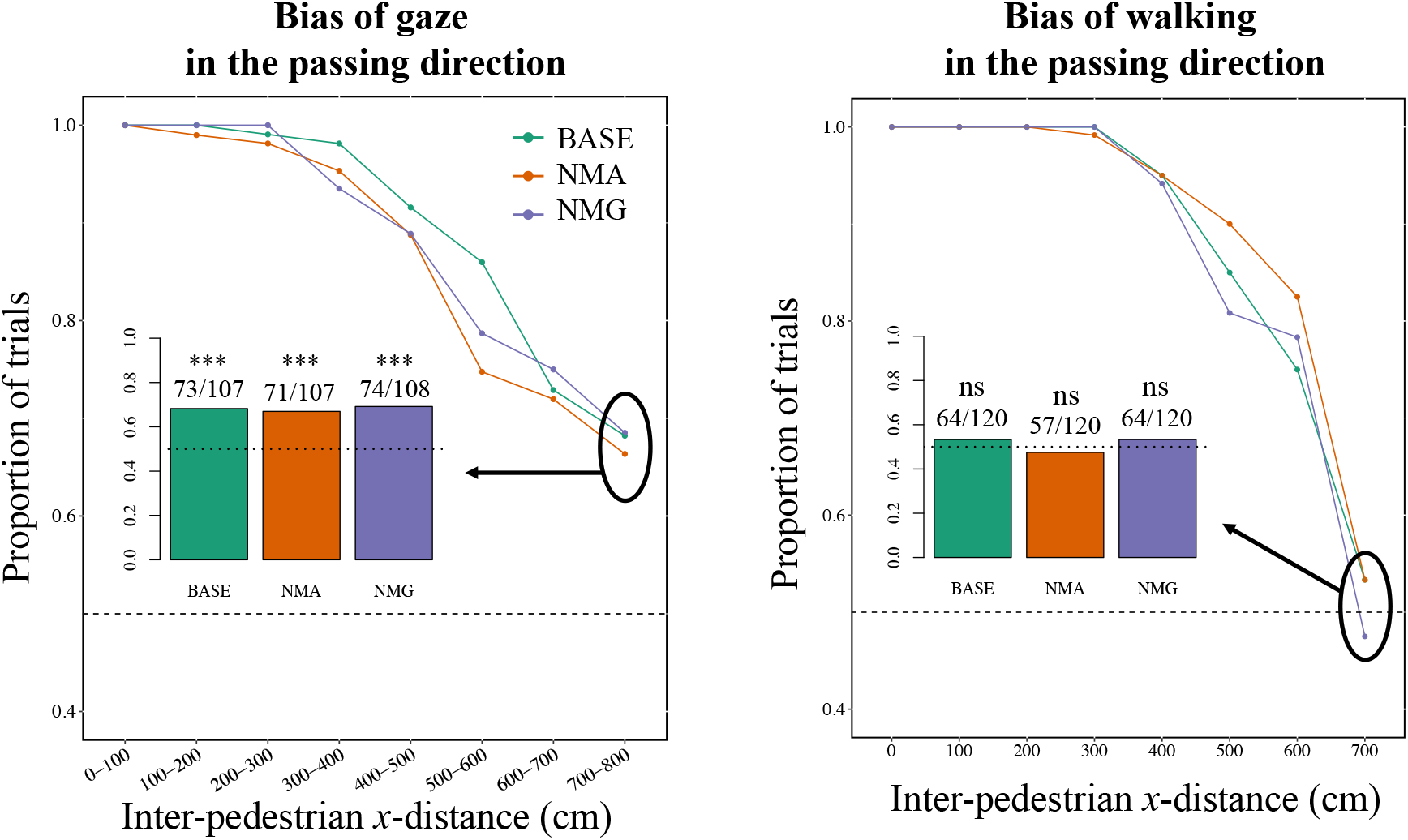
Relation between directions of walking and gaze. Proportion of trials in which participants looked more in the ultimate passing direction during sections of inter-pedestrian distance along the *x*-axis (left). Proportion of trials in which participants were on the side of passing at a certain inter-pedestrian distance (700, 600, … 0 cm) on the basis of the *y*-position at the maximum distance (800 cm) (right). Black dashed lines represent a 0.5 (chance) level. Insets show the details of the proportions immediately after trial onset (left: during 700–800 cm; right: at 700 cm). Asterisks indicate significant departures (***p < 0.001; ns, p > 0.05) from the chance level (dashed lines).

## Discussion

We aimed to determine whether pairs of pedestrians coordinated their walking with each other and what role their gaze had during walking. To this end, a combination of motion- and eye-tracking systems was introduced in a simple avoidance experiment. We found that two interacting pedestrians showed joint actions depending on their partner’s motions rather than following an independently predetermined path. As we predicted, distracting the anticipatory ability of pedestrians decreased their coordination while walking. Interestingly, merely blocking one pedestrian’s gaze information had little effect on their coordinated behavior, ruling out the possibility that mutual gaze impacts pedestrian anticipation. Moreover, eye-movement analysis revealed that pedestrians keep their eyes on an oncoming distracted pedestrian for a long time, suggesting that their gaze behavior depends on their uncertainty. In addition, pedestrians looked ahead toward the ultimate passing direction before they began to walk in that direction, suggesting that they use predictive gaze individually rather than mutually to decide which direction each of them should use to pass.

We observed pairs of pedestrians spontaneously perform complex joint-motions when they could mutually anticipate the direction. What functional advantage does this type of continuous coordination provide? When pedestrians are walking toward other pedestrians, it would take too long to accumulate a complete set of information before making the decisions necessary to avoid each other. Rather, they need to continue to update their decision-making as they continuously acquire information while walking [14, 17]. Pedestrians therefore should balance the benefits of making the correct decision (e.g., safely avoiding collisions) with its costs (e.g., the time constraint to reach the destination). On one hand, making a large turn in either direction may enhance collision avoidance, but it would also entail a higher cost because of the longer travel distance. On the other hand, walking in a straight line would minimize energetic cost and time to reach the destination, but may result in a situation where a collision is imminent. Making sharp turns in response to this situation eventually would require active muscular intervention and disturb the preferred gait cycle [31]. Hence, pedestrians must coordinate their motions constantly through real-time negotiation to make a trade-off between the efficiency of the preferred walking path and the need to move laterally to avoid collisions. These coordinated motions will particularly matter in a crowded situation because behavioral coordination among pedestrians helps to smooth pathways through a crowd and facilitate an efficient transition to emergent pattern formation at the collective level [14].

Conversely, distracting one of the two pedestrians to interfere with their ability to anticipate their partner’s motion substantially altered both pedestrians’ behaviors (Fig. 1) and worsened their coordination (Fig. 3). How does this type of experimental manipulation cause such changes even though at least one of the participants was not distracted? Consider the results from the eye-movement analysis. Pedestrians kept their eyes on the distracted partner until they passed by each other (Fig. 4B), even though they had attained sufficient distance to pass safely (Fig. 2A). We speculate that pedestrians were more uncertain about the motions of distracted walkers than they were of non-distracted ones, so that they have to continue to seek information. This idea is consistent with studies on decision-making in sensorimotor locomotion control, according to which people explore the environment with their eyes to navigate and dedicate more of their visual resources on uncertain locations to gain information [30, 31]. This explanation has usually been applied to uncertainty regarding the stability of an environment (e.g., whether terrain is flat [certain] or rocky [uncertain]); however, it could be applied to the uncertainty regarding the dynamics of an environment. From this point of view, we suggest that distracted pedestrians are themselves mobile distractions for other pedestrians.

In line with this, it may be hard to maintain a sufficient distance from these types of mobile distractions in a human crowd due to the restricted amount of free space. In addition, the resources required for visual attention and/or other types of cognitive processing are limited [32]; hence, pedestrians encountering distracted ones could also have difficulty in paying sufficient attention to all of the surrounding pedestrians. These physical and cognitive restrictions could lead to the situation where a collision is imminent and a person must make an immediate turn to avoid the collision. In other words, mobile distractions in a crowd impose indirect restrictions on pedestrians who are not directly distracted. According to Murakami et al. [14], distracting only on few individuals in a crowd increased the occurrence of imminent collisions and disturbed the self-organization process. Indirect restrictions propagated by mobile distractions may provide a possible explanation of this previous result.

Somewhat unexpectedly, having pedestrians wear mirrored sunglasses to block the mutual gaze between them did not influence their walking behavior. We conclude, therefore, that it is not plausible that pedestrians infer other people’s movement trajectories from the gaze direction, as was observed in a previous study [20]. Body motion cues instead are a possible source of visual information, as was noted by Lynch et al. [21], but when does information from body cues become available? We observed that pedestrians looked ahead toward their ultimate passing direction, which may hint at the answer to this question. Although additional data are clearly required, if possible future movement directions of pedestrians are expressed in their eye movements (i.e., looking ahead) before they actually walk in a given direction, that same future information may also be expressed in other body motions, such as movements and/or orientations of the head, shoulder, chest, and so on. In addition, because gaze distributions of one pedestrian crossed all over the body of the other (Figure 4A), it is possible that cues of future motions are embedded in a pattern of whole-body motions [33, 34], rather than in particular parts of the body. It is possible that these body motions are available for pedestrians to anticipate future motions in others before they are executed. Therefore, walkers may implicitly negotiate potential future motions by using body motions.

Overall, we addressed behavioral coordination between mutually anticipating pedestrians and the role of gaze when people pass each other. We expect that our findings will help enhance our understanding of mutual anticipation in other human and animal interactions [35-38], and may prove to be useful in various fields, including research on improvisational motions, pedestrian transportation, and robotic navigation [39].

## Materials and methods

### Experimental design

#### Participants

Twenty male student participants were recruited for the study (mean age ± SD = 22.2 ± 1.06 years). All participants had normal or corrected-to-normal vision. Each was asked to wear a black T-shirt and yellow cap for ease of video analysis; they also wore a white mask to prevent the spread of infectious diseases. Written informed consent was obtained from all participants prior to the start of the study. The study was approved by the Ethics Committee of Kyoto Institute of Technology.

#### Apparatus

The experiment was conducted in the gymnasium of the Kyoto Institute of Technology, Kyoto, on 22 and 23 March 2021. The floor of the study area was covered with a PVC gym floor cover so that the lines on the gym floor were concealed and participants could wear their own outdoor shoes to walk. For the purpose of efficient operation, we prepared two equivalent straight mock-corridors (length = 8 m, width = 1.7 m each). The corridors were constructed with partitions (height = 74 cm) and were 4.73 m from each other. We could thus carry out two experiments in parallel.

#### Task

Before the trial, two participants were instructed to take positions on the middle of the start lines at either end of the corridor, respectively. At the start signal, they were asked to walk in their usual way toward the opposite end.

In addition to observing the pedestrians’ coordinated behavior, we attempted to observe the influence of mutual anticipation and mutual gaze. We therefore set three experimental conditions: baseline (BASE), no mutual anticipation (NMA), and no mutual gaze (NMG).

One of two participants (P1) was asked to do additional tasks while walking in the NMA and NMG conditions; there was no additional task for either participant during the BASE condition. In the NMA condition, P1 was instructed to walk while using a mobile phone. This was intended to distract the pedestrians visually and potentially intervene in anticipatory interactions between them. Distracting pedestrians with a mobile phone task made it more difficult for them to anticipate their neighbors’ motions in previous studies [14, 24, 25], indicating that pedestrian motion is strongly influenced by other oncoming pedestrians even when they are well separated [11]. We adopted the mobile phone task used in the pedestrian experiment of Murakami et al. [14]; participants were asked to do a simple arithmetic task (a single-digit addition problem) on their mobile phones. Specifically, P1 participants were asked to start solving problems from the moment they heard the start signal until they exited at the opposite end of the mock corridor and to endeavor to solve as many problems as possible. In the NMG condition, P1s were instructed to walk wearing a one-way mirrored sunglasses (FILA, visible light transmission = 15%) (Figure S6) to block gaze information from P1 to P2; P2 wore eye-tracking glasses during this experiment (Tobii Glasses Pro 3; Figure S6).

Before starting the experiments, we informed all participants that there could be oncoming walkers who would walk while using a mobile phone or wearing sunglasses and explained how to perform the mobile phone tasks. We did this to ensure that all of the participants knew of the existence of manipulated pedestrians during the experiment (i.e., to avoid a sort of progressive learning process in which participants gradually begin to understand details of the experimental procedure). We conducted two pretest trials under the BASE condition to confirm participants had correctly understood instructions given to them before conducting the main experiments.

#### Procedure

The 20 participants were divided into four groups of five people each. Two sessions were performed each day, for a total of four independent sessions. Each session was carried out with one of the four groups and was divided into five blocks, with each block having three subblocks, and each subblock was further divided into three trials (3 trials × 3 subblocks × 5 blocks × 2 corridors = 90 trials in each session). In each block, two participants were assigned to wear the eye-tracking glasses (i.e., they were the P2 participants), and two of the other three participants were assigned to do additional tasks according to the experimental condition (i.e., P1 participants), one in each experimental corridor. The remaining participant in each group did not actively participate in the subblock and was allowed to rest. P2 participants remained in the same role during a single block, but P1 members were shuffled between the three subblocks so that each P2 walked with each of the P1s in a single block. In each subblock, three trials (one for each experimental condition; i.e., BASE, NMA, and NMG) were conducted in a randomized order. Participants were shuffled among the five blocks so that each one was assigned to the P2 role in two blocks (18 trials) and the P1 role in 18 trials. In total, the experiments were replicated 120 times under each condition (4 sessions × 30 trials per session per condition).

#### Post-experiment questionnaire

To measure the extent to which participants felt that the oncoming person paid attention to them, a questionnaire was administered after all experiments were performed. Participants were asked to respond to the following item (translated from Japanese): “the oncoming person paid attention to me while I was walking with wearing eye-tracking glasses.” They indicated their response on a visual analog scale ranging from –3 (completely disagree) to +3 (completely agree) for each of the three experimental conditions (i.e., there were three questions in total). The questionnaire items were presented in a randomized order.

### Body movement tracking and analysis

The experiments were recorded from above with GoPro Hero 8 camcorders equipped with a linear digital lens (2.7K, 60 fps); the cameras were fixed at a height of 10 m above the floor. From the video images, we tracked the time series of each individual’s positions frame-by-frame by using the image-processing software PeTrack [40].

#### Normalizing the time axis

Since the time it took for the pedestrians to pass each other varied for each trial, the trials each had different numbers of data points. Therefore, we used an average curve to reflect the shape of the individual curves in the analysis (Figure 3A). To calculate the average curve, individual curves were normalized so that a relative time axis ranging from 0 (trial onset) to 1 (the moment of passing). The data were then graphically re-digitized at an arbitrarily high spatial frequency so that all curves had 216 data points [41, 42]; after this transformation, the values (i.e., the differences between the indicators) at each point were averaged.

### Eye-movement tracking and analysis

To investigate where and when participants looked during the experiment, we asked P2 participants to wear an eye-tracking system (Tobii Pro Glasses 3, 100 Hz) mounted with a scene camera (HD, 25 fps) facing outward; the system was calibrated with the help of the experimenter. Before the main experiments were conducted, each participant took part in a familiarization session where they walked while wearing the experimental apparatus. This eye-tracking system measured the 3D gaze vectors of participants, from which 2D gaze points (in pixels) projected on a video surface (i.e., the far plane of the viewing frustum of the scene camera) were calculated. Recording data were processed, analyzed, and exported by a dedicated software tool (Tobii Pro Lab). We analyzed the gaze behavior of participants from the trial onset (i.e., the start signal) until the image of the oncoming participant’s cap on the video surface was no longer within the scene camera’s angle of view. In 20 of the 360 trials, gaze data were lost because of technical problems with the eye-tracking system. Moreover, when we calculated the mis-acquisition rate of gaze points, the mean rate for one participant was greater than 0.3, even though mean rate for all trials was 0.076 ± 0.109; all gaze data acquired from this participant were excluded (18 trials). The gaze data of the remaining 322 trials acquired from 19 participants were used in the analysis.

To investigate how gaze points are distributed around the images of oncoming pedestrians (i.e., P1) on the video surface, we tracked P1 head (cap) positions on the video surface in pixels by using image-processing software (Library Move-tr/2D) and calculated the gaze points relative to P1’s head positions on the video surface. However, because there is no information about physical dimensions in the video surface, we converted these gaze points determined from pixel-coordinates to cm-coordinates by using the camera parameters (camera resolution and angle of view) and the distance between P1 and P2 calculated from the walking trajectories at each point in time. Mathematically, each component of the cm-coordinates of gaze points relative to P1’s head position at time *t*, (*x*_*g*_(*t*), *y*_*g*_(*t*)), was calculated as

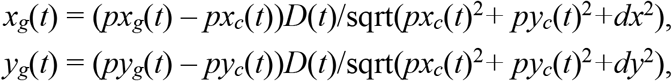

where *px*_*g*_(*t*) and *py*_*g*_(*t*) represent the horizonal and vertical components of pixel-coordinates of a gaze point on the video surface at time *t*, respectively; *D*(*t*) is the distance between P1 and P2 at time *t*; *px*_*c*_(*t*) and *py*_*c*_(*t*) represent the horizonal and vertical components of pixel-coordinates of P1’s head position on the video surface at time *t*, respectively; and *dx* and *dy* are constant values calculated from the scene camera’s parameters as

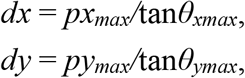

where *px*_*max*_ = 960 (pixel), *py*_*max*_ = 540 (pixel), *θ*_*xmax*_ = 0.829 (rad), and *θ*_*ymax*_ = 0.549 (rad). Note that the origin of the pixel-coordinates was set to the center of the video surface. In this way, we calculated how far P2’s gaze points are from P1’s head positions on a plane that is parallel to the video surface and includes P1’s head position at each point in time (see Fig. S7).

#### Calculating the proportion of gaze points at the AOI

We calculated the proportion at which each pedestrian gazed at the AOI in each section of the inter-pedestrian distance (Figure 4B). We then averaged the proportions for each participant under each condition and treated them as a feature of each participant. We then compared within-participant means between BASE and each of the other conditions.

### Synchronization of the timestamps between the eye and body movements tracking systems

To synchronize the timestamps between the overhead video cameras for body-movement tracking and the scene camera mounted on the eye-tracking glasses, a laptop computer (MacBook Pro, refresh rate = 60 Hz) displaying a switch from a white full screen image to a black one was recorded by all cameras together each time after the wearers of the eye-tracking glasses were changed (i.e., before each block started). We used a frame where the black full-screen image was completely displayed as the timestamp for synchronization.

### Correspondence among different recording rate data

The recording rates differed among the eye-tracker (100 Hz), its scene camera (25 Hz), and the overhead videos (60 Hz) for body-movement tracking. We corresponded each gaze point with a frame of the overhead camera with a timestamp that was the nearest to that of the gaze point and with a frame of the scene camera in the same manner.

## Statistical analysis

Welch’s t tests were used to test differences in the distance between two pedestrians at the moment of passing, the time taken from the trial onset to the moment of passing, and mean differences between two pedestrians in the angular deviation and change in speed. Paired t tests were conducted to test differences in the proportion of gaze points on the AOI and questionnaire ratings. When making multiple comparisons, p values were adjusted by using false discovery rate control [43]. Additionally, we performed exact binomial tests to compare the number of trials in which participants moved in the ultimate passing direction from the start position and those in which participants looked more in the ultimate passing direction. Details of the statistics are reported in Tables S1–3. All statistical analyses were conducted using R version 3.6.0 (The R Foundation for Statistical Computing, Vienna, Austria).

## Notes

### Competing Interest Statement

The authors have declared no competing interest.

