## Supplemental Information for "Spontaneous behavioral coordination between pedestrians emerges through mutual anticipation rather than mutual gaze"

This SI contains figure S1-S7 and table S1-S3.

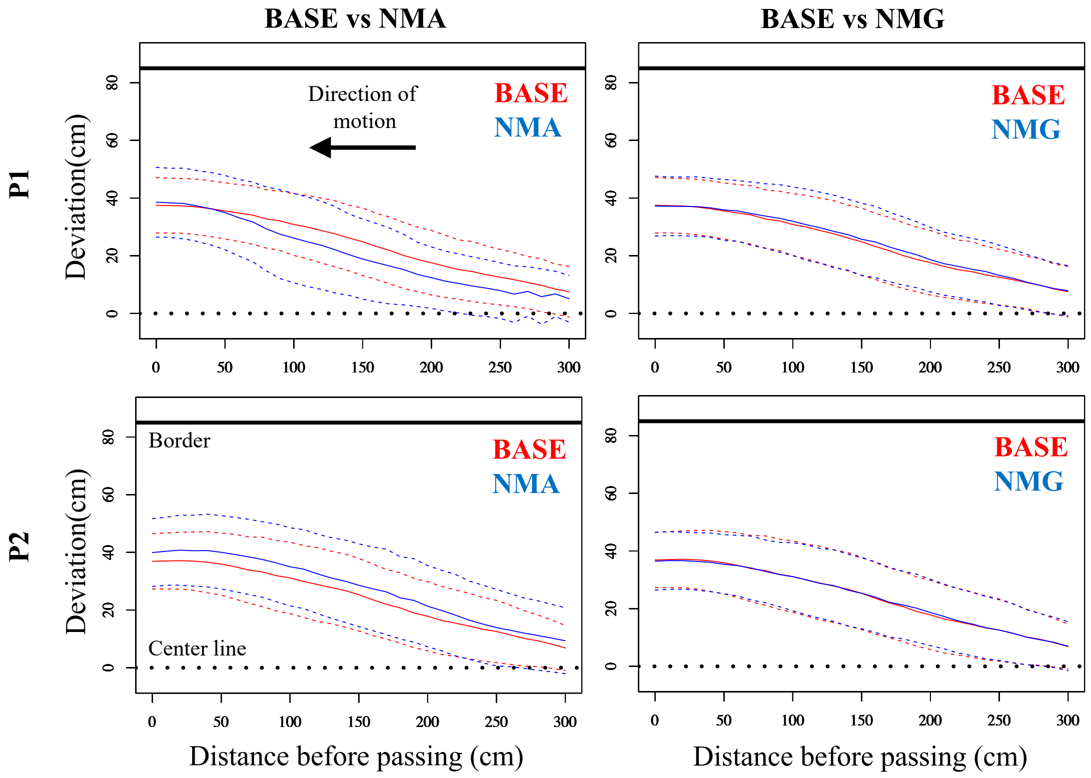


**Figure S1. Comparisons of trajectories between the BASE and other conditions.** Solid lines show average lateral deviation before passing under each condition. Dashed lines represent the standard deviation.

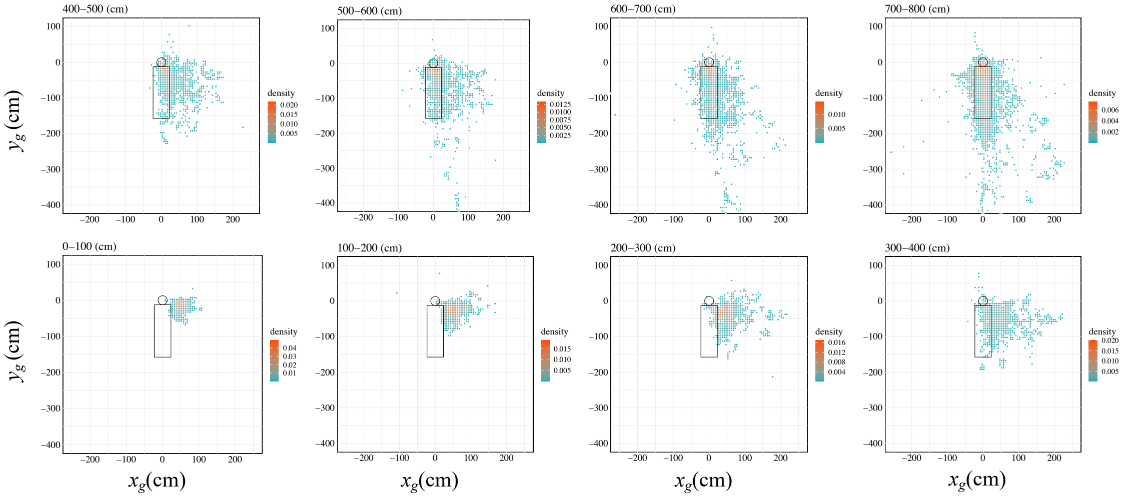


**Figure S2. Gaze behavior around oncoming pedestrians while walking under the BASE condition.** Two-dimensional distributions of gaze points around oncoming pedestrians during sections of distance along the *x*-axis between two pedestrians. $x_g$ and $y_g$ represent the horizonal and vertical components of the cm-coordinates of gaze points relative to P1's head position, respectively (see the main text). The human-like silhouette composed of a circle and a rectangle is an area of interest representing the oncoming pedestrian's body area. Passing direction was positioned to be right. Bins are 5 × 5 (cm).

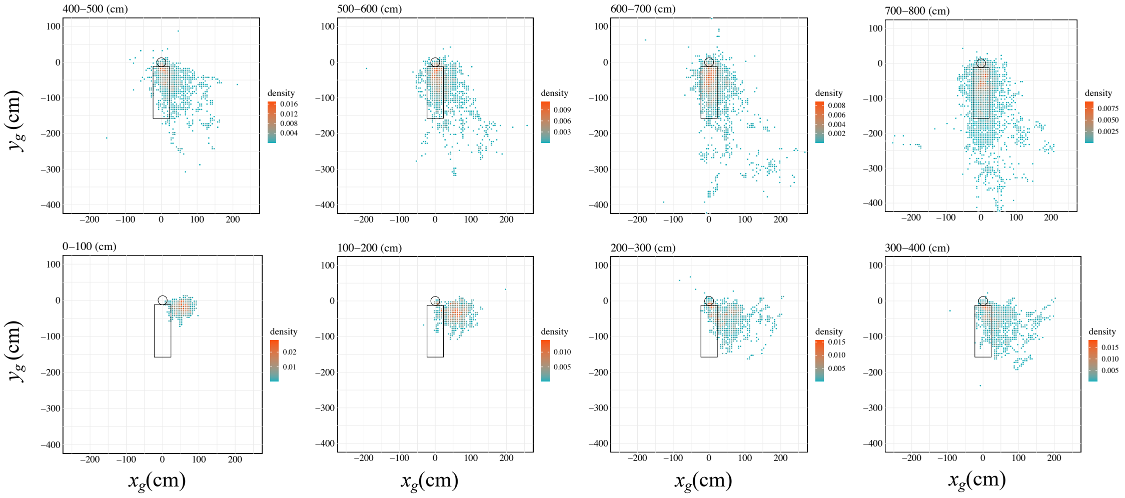


**Figure S3. Gaze behavior around oncoming pedestrians while walking under the NMA condition.** See Figure S2 for a description of the two-dimensional distributions of gaze points.

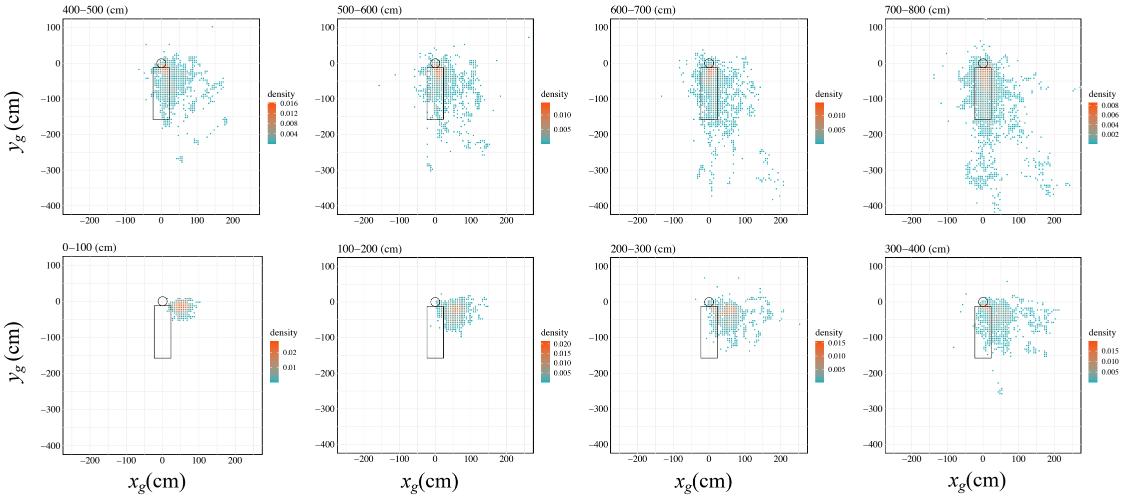


**Figure S4. Gaze behavior around oncoming pedestrians while walking under the NMG condition.** See Figure S2 for a description of the two-dimensional distributions of gaze points.

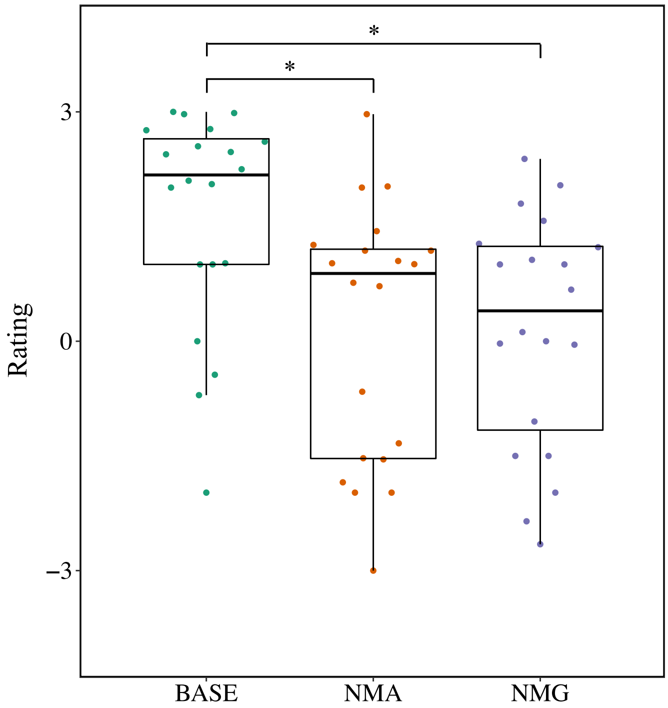


**Figure S5. Results of the post-experiment questionnaire survey.** We asked participants (P2) to answer three items for each of the three different experimental conditions concerning how they perceived that the oncoming pedestrian (P1) paid attention to them. Each data point represents a participant (on a visual analog scale of −3 [completely disagree] to 3 [completely agree]). Box-and-whisker plots represent the median (central thick line), first and third quartiles (box), and 1.5× the interquartile range of the median (whiskers). Asterisks indicate statistical significance in a comparison between the BASE condition and each of the other conditions (*$p < 0.05$).

(Fig.S6)

**Figure S6. Sunglasses, smart phone, and eye-tracking glasses used in the present experiment.**

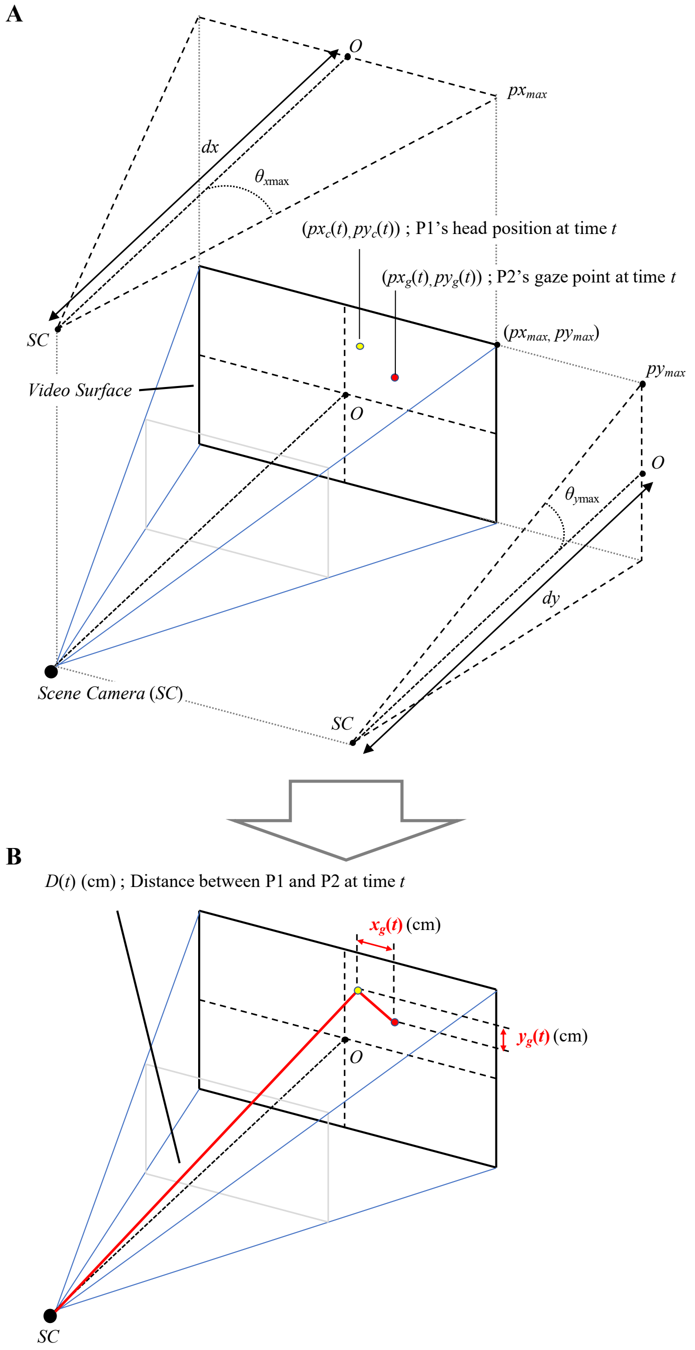


**Figure S7. Diagrams of head positions, gaze points, and camera positions used in the analyses of eye-movement data.** We obtained the cm-coordinates of gaze points relative to P1's head position ($x_g(t)$, $y_g(t)$) (**B**) from the pixel-coordinates of the gaze point ($px_g(t)$, $py_g(t)$) and P1's head position ($px_c(t)$, $py_c(t)$) by using the camera parameters ($px_{max}$, $py_{max}$, $\theta_{xmax}$, and $\theta_{ymax}$) and the distance between P1 and P2 (D($t$)) (**A**). *O* represents the coordinates' origin. See the Materials and Methods section in the main text for more details.

**Table S1. Summary of Welch's t test used in the study.**

| Measure | Comparison | t | p value | Cohen's d |
|---|---|---|---|---|
| Distance when passing | BASE (n = 120) vs. NMA (n = 120) | −3.09 | 0.004 | −0.39 |
| Distance when passing | BASE (n = 120) vs. NMG (n = 120) | 0.43 | 0.66 | 0.0056 |
| Time until passing | BASE (n = 120) vs. NMA (n = 120) | −6.93 | <0.001 | −0.89 |
| Time until passing | BASE (n = 120) vs. NMG (n = 120) | −0.188 | 0.85 | −0.024 |
| Difference of angular deviation; BASE | Actual pairs (n = 120) vs. random pairs (n = 120) | −3.66 | <0.001 | −0.47 |
| Difference of angular deviation; NMA | Actual pairs (n = 120) vs. random pairs (n = 120) | −2.95 | 0.0033 | −0.38 |
| Difference of angular deviation; NMG | Actual pairs (n = 120) vs. random pairs (n = 120) | −3.58 | <0.001 | −0.46 |
| Difference of change in speed; BASE | Actual pairs (n = 120) vs. random pairs (n = 120) | −4.92 | <0.001 | −0.63 |
| Difference of change in speed; NMA | Actual pairs (n = 120) vs. random pairs (n = 120) | −3.1 | 0.002 | −0.4 |
| Difference of change in speed; NMG | Actual pairs (n = 120) vs. random pairs (n = 120) | −5.45 | <0.001 | −0.69 |

**Table S2. Summary of within-subjects paired t tests used in the study.**

| Measure | Comparison | t | p value | Cohen's d |
|---|---|---|---|---|
| Proportion of gaze on AOI; 700–800 cm | BASE (n = 19) vs. NMA (n = 19) | 0.06 | 0.951 | 0.01 |
| Proportion of gaze on AOI; 700–800 cm | BASE (n = 19) vs. NMG (n = 19) | 0.34 | 0.951 | 0.07 |
| Proportion of gaze on AOI; 600–700 cm | BASE (n = 19) vs. NMA (n = 19) | -2.7 | 0.015 | -0.31 |
| Proportion of gaze on AOI; 600–700 cm | BASE (n = 19) vs. NMG (n = 19) | -2.91 | 0.015 | -0.33 |
| Proportion of gaze on AOI; 500–600 cm | BASE (n = 19) vs. NMA (n = 19) | -2.37 | 0.058 | -0.31 |
| Proportion of gaze on AOI; 500–600 cm | BASE (n = 19) vs. NMG (n = 19) | -0.99 | 0.335 | -0.14 |
| Proportion of gaze on AOI; 400–500 cm | BASE (n = 19) vs. NMA (n = 19) | -1.6 | 0.256 | -0.25 |
| Proportion of gaze on AOI; 400–500 cm | BASE (n = 19) vs. NMG (n = 19) | -0.28 | 0.78 | -0.04 |
| Proportion of gaze on AOI; 300–400 cm | BASE (n = 19) vs. NMA (n = 19) | -2.49 | 0.046 | -0.42 |
| Proportion of gaze on AOI; 300–400 cm | BASE (n = 19) vs. NMG (n = 19) | -0.64 | 0.528 | -0.09 |
| Proportion of gaze on AOI; 200–300 cm | BASE (n = 19) vs. NMA (n = 19) | -3.37 | 0.007 | -0.63 |
| Proportion of gaze on AOI; 200–300 cm | BASE (n = 19) vs. NMG (n = 19) | -1.21 | 0.242 | -0.24 |
| Proportion of gaze on AOI; 100–200 cm | BASE (n = 19) vs. NMA (n = 19) | -2.99 | 0.016 | -0.48 |
| Proportion of gaze on AOI; 100–200 cm | BASE (n = 19) vs. NMG (n = 19) | 0.01 | 0.992 | 0 |
| Proportion of gaze on AOI; 0–100 cm | BASE (n = 18) vs. NMA (n = 18) | -0.55 | 0.989 | -0.09 |
| Proportion of gaze on AOI; 0–100 cm | BASE (n = 18) vs. NMG (n = 18) | −0.13 | 0.99 | −0.02 |
| Rating of questionnaire | BASE (n = 20) vs. NMA (n = 20) | 2.75 | 0.011 | 0.96 |
| Rating of questionnaire | BASE (n = 20) vs. NMG (n = 20) | 3.12 | 0.012 | 1.01 |

**Table S3. Summary of binominal tests used in the study.**

| | Distance or distance section (cm) | Condition | Trial toward passing direction | Total | p value |
|---|---|---|---|---|---|
| Results of body movement | 700 | BASE | 64 | 120 | 0.52 |
| | 700 | NMA | 64 | 120 | 0.52 |
| | 700 | NMG | 57 | 120 | 0.64 |
| | 600 | BASE | 90 | 120 | <0.001 |
| | 600 | NMA | 99 | 120 | <0.001 |
| | 600 | NMG | 94 | 120 | <0.001 |
| | 500 | BASE | 102 | 120 | <0.001 |
| | 500 | NMA | 108 | 120 | <0.001 |
| | 500 | NMG | 97 | 120 | <0.001 |
| | 400 | BASE | 114 | 120 | <0.001 |
| | 400 | NMA | 114 | 120 | <0.001 |
| | 400 | NMG | 113 | 120 | <0.001 |
| | 300 | BASE | 120 | 120 | <0.001 |
| | 300 | NMA | 119 | 120 | <0.001 |
| | 300 | NMG | 120 | 120 | <0.001 |
| | 200 | BASE | 120 | 120 | <0.001 |
| | 200 | NMA | 120 | 120 | <0.001 |
| | 200 | NMG | 120 | 120 | <0.001 |
| | 100 | BASE | 120 | 120 | <0.001 |
| | 100 | NMA | 120 | 120 | <0.001 |
| | 100 | NMG | 120 | 120 | <0.001 |
| Results of eye movement | 800–700 | BASE | 73 | 107 | <0.001 |
| | 800–700 | NMA | 71 | 107 | <0.001 |
| | 800–700 | NMG | 74 | 108 | <0.001 |
| | 700–600 | BASE | 78 | 107 | <0.001 |
| | 700–600 | NMA | 77 | 107 | <0.001 |
| | 700–600 | NMG | 81 | 108 | <0.001 |
| | 600–500 | BASE | 92 | 107 | <0.001 |
| | 600–500 | NMA | 80 | 107 | <0.001 |
| | 600–500 | NMG | 85 | 108 | <0.001 |
| | 500–400 | BASE | 98 | 107 | <0.001 |

| | | | | | |
|---|---|---|---|---|---|
| | 500–400 | NMA | 95 | 107 | <0.001 |
| | 500–400 | NMG | 96 | 108 | <0.001 |
| | 400–300 | BASE | 105 | 107 | <0.001 |
| | 400–300 | NMA | 102 | 107 | <0.001 |
| | 400–300 | NMG | 101 | 108 | <0.001 |
| | 300–200 | BASE | 105 | 106 | <0.001 |
| | 300–200 | NMA | 105 | 107 | <0.001 |
| | 300–200 | NMG | 108 | 108 | <0.001 |
| | 200–100 | BASE | 102 | 102 | <0.001 |
| | 200–100 | NMA | 98 | 99 | <0.001 |
| | 200–100 | NMG | 104 | 104 | <0.001 |
| | 100–0 | BASE | 93 | 93 | <0.001 |
| | 100–0 | NMA | 90 | 90 | <0.001 |
| | 100–0 | NMG | 94 | 94 | <0.001 |
